# Hi-TOM: a platform for high-throughput tracking of mutations induced by CRISPR/Cas systems

**DOI:** 10.1101/235903

**Authors:** Qing Liu, Chun Wang, Xiaozhen Jiao, Huawei Zhang, Lili Song, Yanxin Li, Caixia Gao, Kejian Wang

## Abstract

The CRISPR/Cas system has been extensively applied to make precise genetic modifications in various organisms. Despite its importance and widespread use, large-scale mutation screening remains time-consuming, labour-intensive and costly. Here, we describe a cheap, practicable and high-throughput screening strategy that allows parallel screening of 96 × N (N denotes the number of targets) genome-modified sites. The strategy simplified and streamlined the process of next-generation sequencing (NGS) library construction by fixing the bridge sequences and barcoding primers. We also developed Hi-TOM (available at http://www.hi-tom.net/hi-tom/), an online tool to track the mutations with precise percentage. Analysis of the samples from rice, hexaploid wheat and human cells reveals that the Hi-TOM tool has high reliability and sensitivity in tracking various mutations, especially complex chimeric mutations that frequently induced by genome editing. Hi-TOM does not require specially design of barcode primers, cumbersome parameter configuration or additional data analysis. Thus, the streamlined NGS library construction and comprehensive result output make Hi-TOM particularly suitable for high-throughput identification of all types of mutations induced by CRISPR/Cas systems.

## Introduction

CRISPR/Cas has emerged as powerful genome editing system and has been widely applied in various organisms (Wang et al. 2014; Zhang et al. 2014; Rodríguez-Leal et al. 2017). However, with the relatively high efficiency and broad applicability of genome editing in different organisms, screening mutations in a large number of samples remains a time-consuming, labour-intensive and expensive process. To obtain sequence mutations induced by genome editing, the traditional method used is to clone PCR-amplified target regions, followed by monoclonal sequencing. Alternatively, simple strategies were also developed to identify mutations by directly analyzing the Sanger sequencing results (Brinkman et al. 2014; Xie et al. 2017). However, for a large number of samples, Sanger sequencing is expensive and the decoding process is time-consuming. In addition, complex chimeric mutations, which are frequently generated by genome editing, remain difficult to decode.

Due to the tremendous progress in terms of speed, throughput and cost, next-generation sequencing (NGS) has been used increasingly in biological research (Goodwin et al. 2016). With NGS, thousands to millions of sequencing reactions can be performed in parallel, which generates a valuable amount of sequence information (Metzker 2010). By taking advantages of NGS technology, high-throughput screening of mutations can be accomplished. Although many tools have been developed to analyze the sequencing data (Guell et al. 2014; Xue and Tsai 2015; Lindsay et al. 2016; Park et al. 2016; Pinello et al. 2016), it remains difficult for researchers who are not familiar with bioinformatics to construct the NGS library and analyze the sequencing data, which constrains the wide application of NGS in labs that conducting genome editing. Here, we developed a simplified and cheap strategy that combines common experiment kits for NGS library construction and a user-friendly web tool for high-throughput analysis of multiple samples at multiple target sites. The work will facilitate the application of NGS sequencing in mutation identification, particularly for laboratories that do not have researchers with NGS or bioinformatics skills.

## Results

To establish a simplified and streamlined workflow to screen mutations induced by CRISPR/Cas systems, we first modified the PCR-based library construction strategy. Library construction included two rounds of PCR, i.e. target-specific and barcoding PCR. The initial PCR primers included a target-specific sequence with common bridging sequences (5’-ggagtgagtacggtgtgc-3’ and 5’-gagttggatgctggatgg-3’) added at the 5’ end (Fig.1A). To ensure the sequencing quality of the mutated sequence, the targeted sites were designed within 10-100 nt of either the forward or reverse target-specific primers (Fig.1A). The annealing temperature of the first PCR was determined by the Tm value of the target-specific sequences. The first-round PCR products were further barcoded during the second-round PCR. Common primers for the second-round PCR include platform-specific adaptor sequence, fixed barcode sequence and bridging sequence (Fig.1B, Supplementary Table 2). To improve the recognition ability and reduce the sample confusion, 4 base barcode that contains at least two nucleotides difference with one another was designed. In addition, to ensure high sequencing quality of barcode, 4 spacer bases were also added between the sequencing primers and barcodes. To establish a standard platform for data analysis, 12 forward and 8 reverse primers were fixed as common primers for the second PCR step. Thus, it was possible to create a uniquely barcoded amplicon for up to 96 samples (Fig.1B, Supplementary Table 1, Supplementary Table 2). For economy, each long primer used in the second PCR step was divided into two short primers, which were mixed in 1:500 proportions in each reaction mix. For convenience, the primers and PCR mix can be pre-assembled as standard 96-well PCR kits (Fig.1B, Supplementary Table 1), so there is no need to prepare PCR mix each time. After the second-round PCR amplification, the products of all samples were mixed with equal amount. If multiple targets (number: N) are required for identification, all products (96 × N) can be mixed as a single sample and sent for NGS company. In most cases, 1 Gb sequencing data, the size of which is about 100 Mb after gzip compression, is enough for mutation analysis.

**Figure 1.**
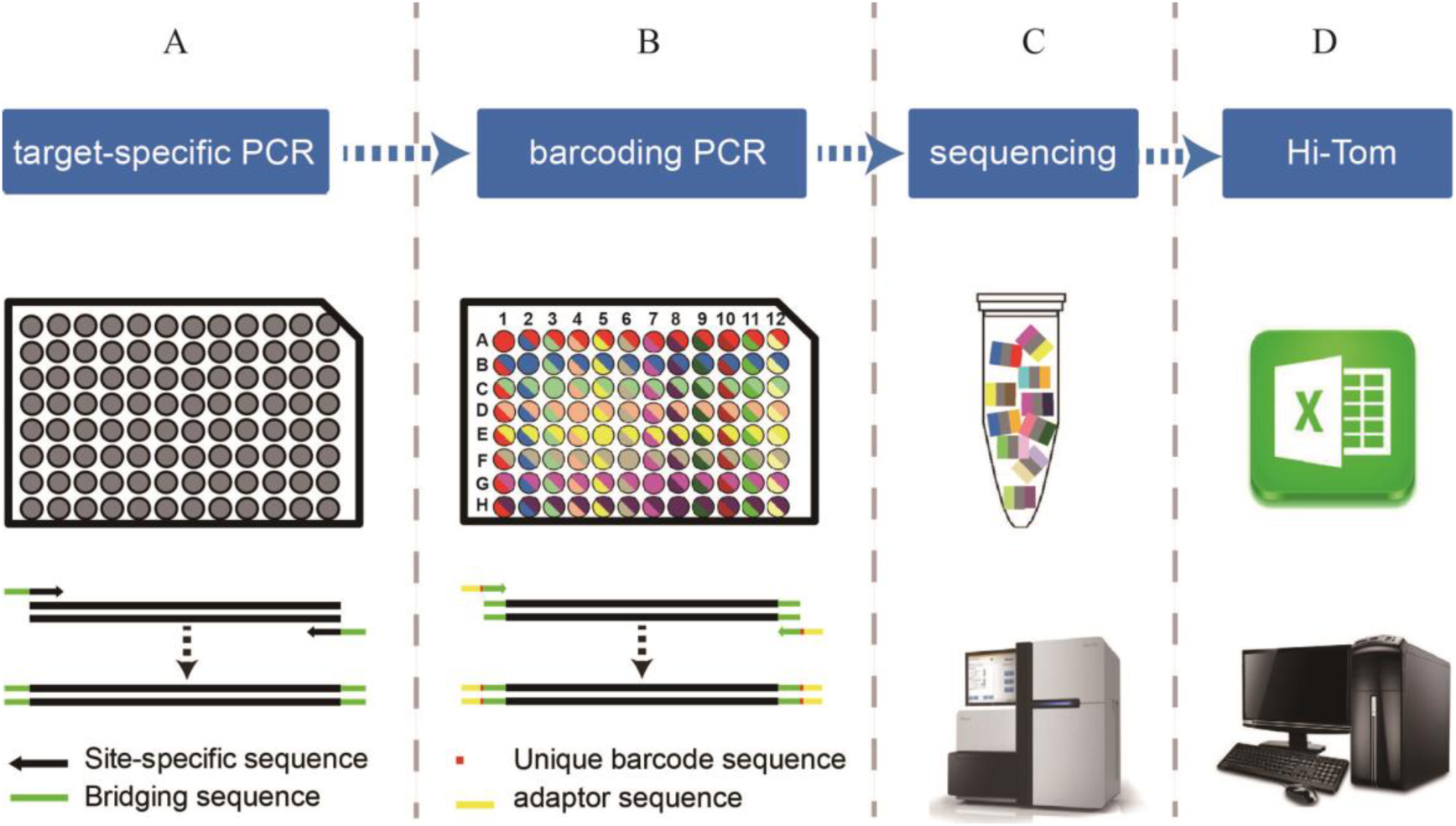
Schematic illustration of the workflow of Hi-TOM. **(A)** The samples are amplified using site-specific primers (target-specific PCR). **(B)** The products of the first-round PCR are used as templates for the second-round PCR (barcoding PCR) in the 96-hole plate kit. By barcoding PCR, the products of each sample are barcoded. **(C)** All products of the second-round PCR are pooled in equimolar amounts in a single tube and sent for NGS. **(D)** Hi-TOM analyses the data sample-by-sample and exports the results in Excel format.

Then we developed a corresponding Hi-TOM platform for high-throughput mutation sequence decoding (http://www.hi-tom.net/hi-tom/). The algorithm behind Hi-TOM was implemented in a Perl script (v5.16.3). The format of the clean data directly produced by NGS instrument or provided by an NGS company is typically gzip-compressed. Hi-TOM uses clean NGS data directly as an input to generate a detailed genome editing report. The first step is entering a working directory (note that filenames cannot include spaces) to avoid conflicts with other tasks. The second step is clicking the ‘Choose File’ buttons to upload the forward reads (named_1.fq.gz) and reverse reads (named_2.fq.gz) (Fig. 1D, Fig. 2A). The third step is to upload the reference sequence length in approximately 1000 nt (typically 500 nt before and after the target sequence) (Fig. 2A). If multiple targets are analyzed simultaneously, the reference sequences of different targets are sorted in a text file in FASTA format and submitted (Fig. 2A). Then, the submit button is clicked and data are uploaded to the server automatically and directly input into the analysis process. And analysis is performed sample-by-sample in an automated manner. When the analysis is completed, a new interface is displayed. The user can then click the ‘download’ button to download the results.

**Figure 2.**
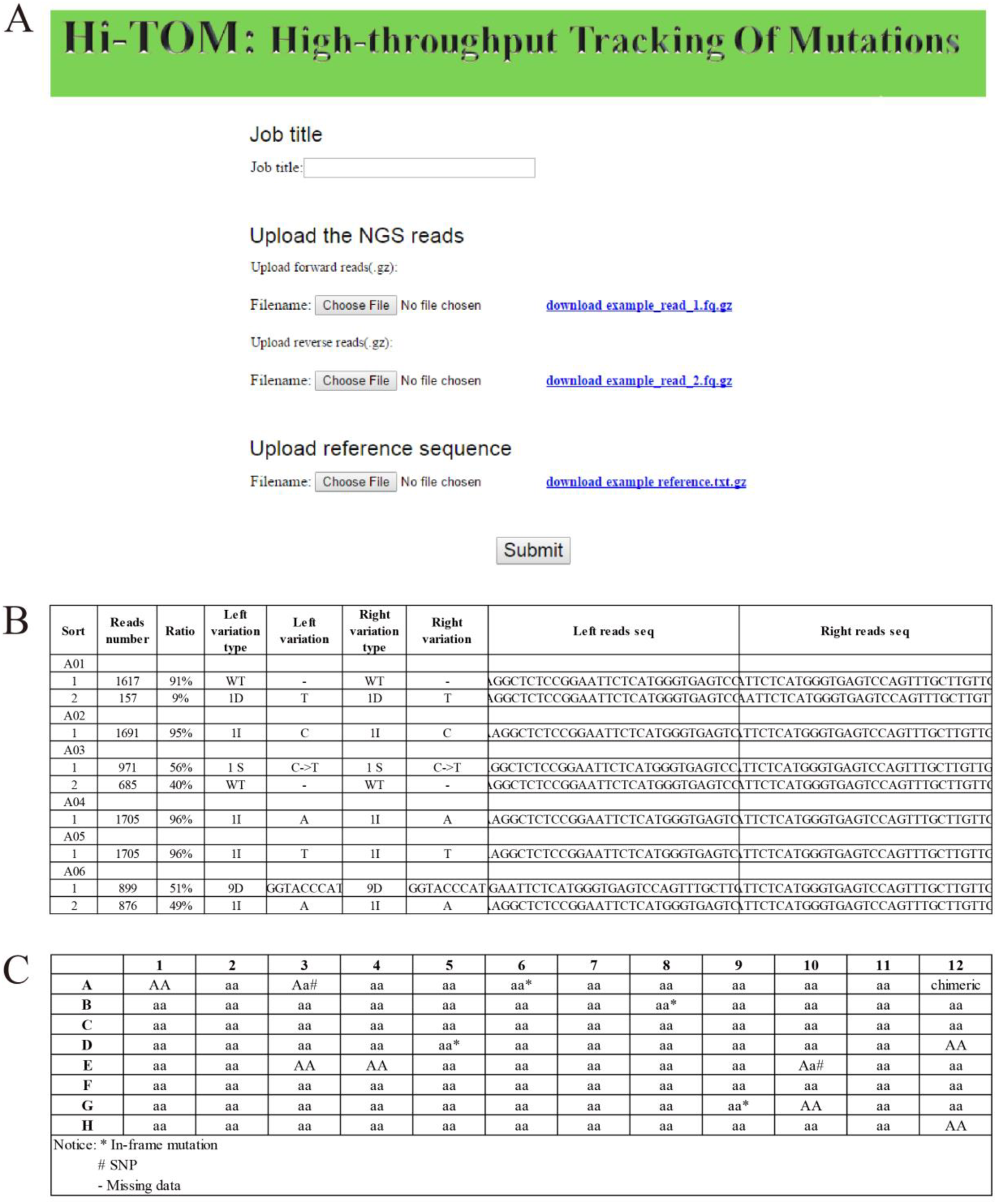
The home page of Hi-TOM and result of example. **(A)** The home page of Hi-TOM. Job title stands for working directory; upload the NGS reads stands for uploading the forward reads (named_1.fq.gz) and reverse reads (named_2.fq.gz); **(B)** An example of Hi-TOM sequence results. The results are summarised in the table. The results are populated in nine columns as follows: sample code and mutant name; read number; ratio; left variation type; left variation; right variation type; right variation; left read sequence; right read sequence. In the variation type volume, I, D and S indicate the insertion, deletion and SNP, respectively. For example, 3D represents the deletion of three bases and 1I represents insertion of one base. In the reads sequence column, the location of the mutation is indicated in lowercase. **(C)** An example of Hi-TOM genotype results. AA stands for wild-type genotype; Aa stands for heterozygous genotype; aa stands for homozygous mutant genotype; * stands for in-frame mutation; # stands for SNP; - stands for missing data.

Hi-TOM analyses the uploaded data and performs quantitative analysis of mutations in three steps. (i) Hi-TOM first finds the reference sequence, converts it to FASTA format, then indexes the reference sequence using BWA software and prepares for the following mapping (Li and Durbin 2009). (ii) The corresponding uploaded data file is then decompressed and the barcodes of each sequence are extracted, the value of each base must be greater than 30 (>Q30). The error sequences resulting from mismatch of primers are eliminated. The sequences are split using the designed barcodes individually. (iii) All reads are mapped to the indexed reference genome after trimming and removing low quality bases using the BWA-MEM (version 0.7.10) algorithm, which shows good performance while mapping reads sequences (Li and Durbin 2009). A Sequence Alignment Map file is then generated. Using this file, the Hi-TOM algorithm screens paired-end reads for sequence alterations, extracts mutation information for each read, and counts the reads number of all type of mutations individually. The aligned results are categorized by mutation type and sorted in descending order of the reads number for each mutation and sample individually. Mutations with the most reads numbers are extracted for each sample, which ensures the decoding of introduced mutations. Hi-TOM integrates the results of 96 samples into a Microsoft Excel document. In each document, the mutation types and positions of each sample are listed in detail, including reads number, ratio, mutation type, mutation bases and DNA sequence (Fig. 2B). As determining the genotype is crucial in many studies, the genotype of each sample is also analyzed and summarized in an additional Excel document (Fig. 2C). Since in-frame mutations and SNP mutations do not necessarily lead to phenotype changes, the samples containing those mutations are highlighted in the table.

In order to demonstrate the applicability and efficiency of the strategy, two sets of genome-edited materials with four targets sites were tested, including rice and human cells. Using the aforementioned strategy, we performed two rounds of PCRs using the site-specific primers and pre-assembled standard 96-well PCR kits. After amplification, the products were pooled, purified and sent for NGS. A total of 1G clean reads were obtained after NGS sequencing. After reads uploading and analyzing, eight excel documents respectively corresponding to those four target sites were automatically generated by Hi-TOM (Fig.2B, Fig.2C, Supplementary Table 3-4). The result shows that sufficient coverage was achieved across all samples at all four target sites (Fig.2B, Fig.2C, Supplementary Table 3-4).

The CRISPR/Cas systems usually generate biallelic, heterozygous and chimeric mutations. The proportions of some mutation types are low and often overlooked. The sensitivity of Hi-TOM was analyzed by comparing with the results of Sanger sequencing. The Sanger sequencing chromatograms showed that the peak values of some mutations are low and complex, which can’t be identified by chromatogram recognition software. In contrast, all these chimeric mutations that contain three, four, even more mutation types were accurately identified by Hi-TOM (Fig.3, Supplementary Table 3-4). In addition to complex chimeric mutations, in-frame mutations and SNP mutations were identified correctly, implying the sensitivity and accuracy of Hi-TOM (Fig.3, Supplementary Table 3-4).

**Figure 3.**
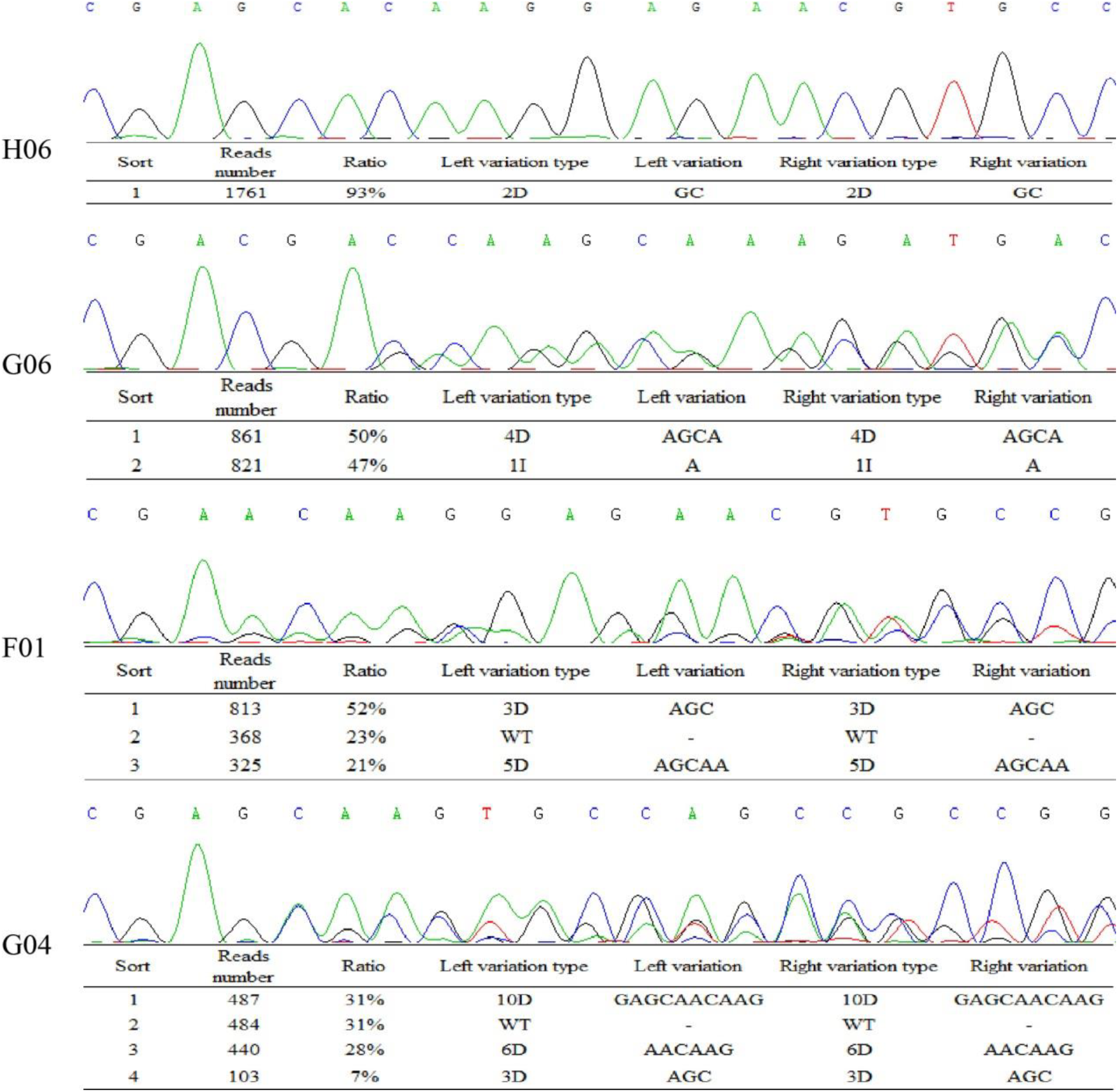
The Sanger sequencing chromatograms and different mutation types tracked by Hi-TOM. H06 represents homozygous mutation; G06 indicates heterozygous mutation; F01 shows chimeric mutation with three variation types; G04 displays chimeric mutation with four variation types.

For polyploid organisms, it still remains a tedious process to analyze mutations on different sets of chromosomes. To test whether Hi-TOM can be used for identification of mutations on different chromosomes in polyploid organism, 64 hexaploid wheat (*Triticum aestivum*, AABBDD) samples were chosen for analysis. To distinguish the A, B and D genomes, Indel and SNP variations on the genome were amplified in the fragment. During Hi-tom analysis, the sequence of A-genome was set as the reference genome. Compared with the amplified sequence of A-genome, the sequence of B-genome has 8bp insertions and 5 SNPs, while the sequence of D-genome has 6bp insertions and 2 SNPs. The results showed that three sets of genomes can be distinguished from each other by analyzing those variations. Meanwhile, the mutations on each genome were successfully tracked (Fig. 4), suggesting the robustness of Hi-TOM in identification of mutations in polyploid organisms.

**Figure 4.**
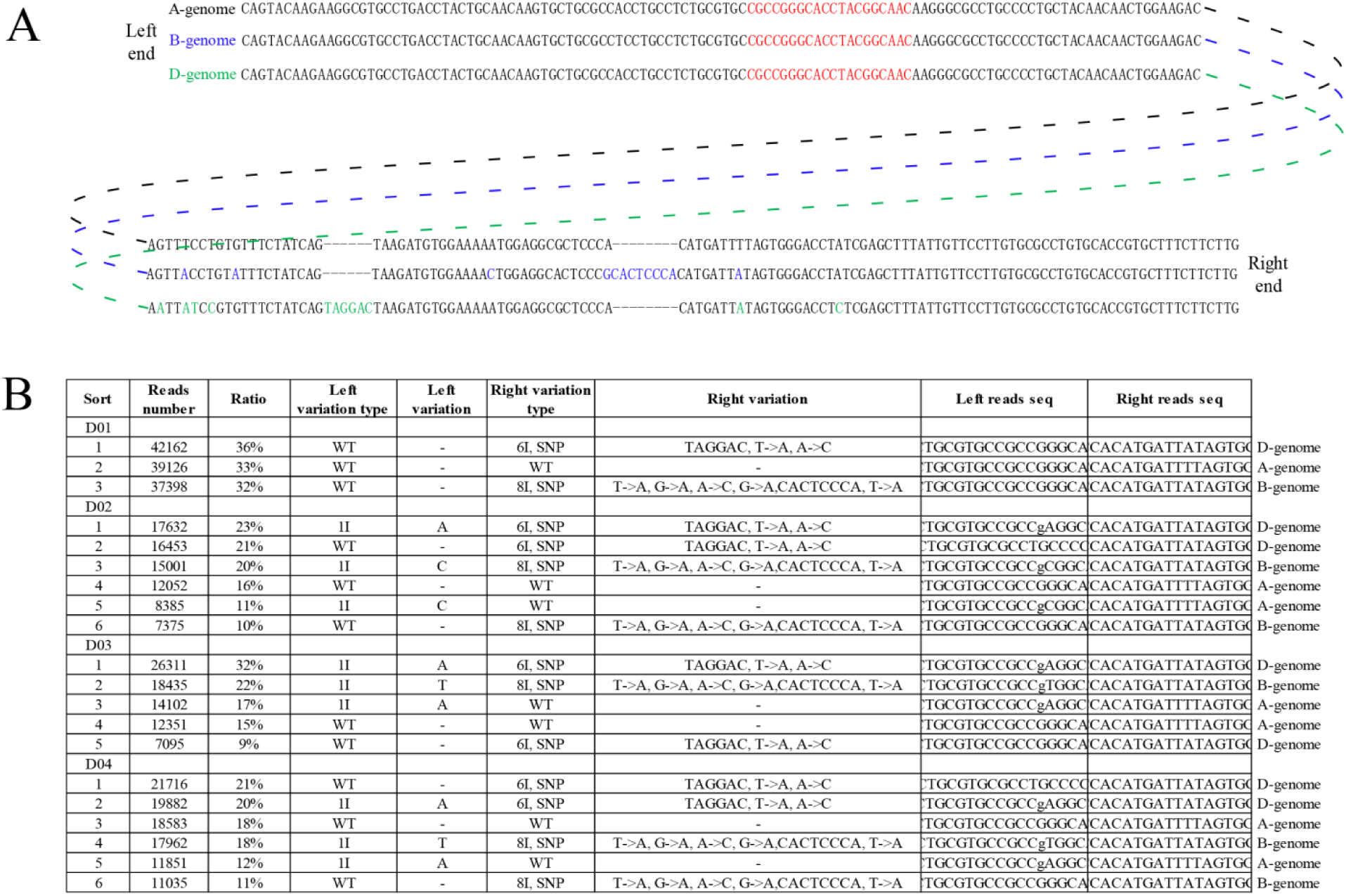
The amplified fragments and Hi-TOM result of hexaploid wheat (*Triticum aestivum*, AABBDD). **(A)** The amplified fragments of A, B and D genomes. The red letters indicate the target sites; The green letters represent sequence variations to distinguish between A genome and B genome; The blue letters show sequence variations to distinguish between A genome and D genome; **(B)** The Hi-TOM result of hexaploid wheat. The mutation variations were displayed in the left variation type while the genomes were shown in the right variation type. WT indicates the A-genome, 8I exhibits the B-genome, and 6I represents the D-genome.

## Discussion

Here we developed a systematic strategy which fixed the second-round PCR primers for the construction of NGS library and established the user-friendly Hi-TOM platform for large-scale mutation identification, which enables the accurate quantification and visualization of mutation outcomes, and comprehensive evaluation of numerous mutation sequences. The possibility of bar coding hundreds of samples with multiple target sites in one sequencing sample makes it competitive economic perspective. As 1 Gb sequencing data is sufficient to analyze hundreds or even thousands of amplicons, the cost can be extremely cheap when analyzing large number of samples or sites. At present, comparing with normal Sanger sequencing, it takes longer time to accomplish the next-generation sequencing. However, with the rapid development of sequencing machine, the sequencing might be accelerated in a very short term. In summary, Hi-TOM provides a high-throughput, economic, sensitive and comprehensive platform for tracking mutations induced by CRISPR/Cas systems.

## Methods

### NGS library construction

The primary PCR was performed to amplify the targeted genomic DNA with a pair of site-specific primers with common bridging sequences (5’-ggagtgagtacggtgtgc-3’ and 5’-gagttggatgctggatgg-3’) added at the 5’ end. The primary amplification was performed in a 20 μL reaction volume containing 50 ng of genomic DNA, 0.3 μM of specific forward and reverse primer, and 10 μL 2×Taq Master Mix (Novoprotein Scientific, China). The secondary amplification was conducted in 20 μL preassembled kits, each containing 10 μL 2×Taq Master Mix, 200 nM 2P-F and 2P-R primer, 2 nM F-(N) and R-(N) primer (Supplementary Table 2), and 1 μL primary PCR product. PCR conditions were 3 min at 94°C (1×), 30 s at 94°C, 30 s at annealing temperature and 30 s at 72°C (33×), followed by 72 °C for 2 min.

### Next generation sequencing

The libraries were sequenced using the Illumina HiSeq platform (Illumina, USA) by the Novogene Bioinformatics Institute, Beijing, China. The concentration of the libraries was initially measured using Qubit^®^2.0 (Life Technologies, USA). The libraries were diluted to 1 ng/μl and an Agilent Bioanalyzer 2100 (Agilent, USA) was used to test the insert size of the libraries. To ensure quality, the SYBR green qRT-PCR protocol was used to accurately dose the effective concentration of the libraries.

### Filtering reads and mapping reads

Paired end (PE) reads with 150 bp were determined and the clean reads were collected from sequenced reads, which were pre-processed to remove adaptors and low quality paired reads. The following criteria were used to remove the low quality reads: i) containing more than 10% ‘N’s; ii) more than 50% bases having low quality value (Phred score <= 5); iii) duplicated reads were removed and coverage values were calculated using SAMTOOLS (Li and Durbin 2009).

## Acknowledgements

We are extremely grateful to Ruiqiang Li from Novogene Bioinformatics Institute for critical reading of the manuscript. We thank Zheng Ruan and his team from Novogene Co., Ltd for NGS technical service. We also thank Yangwen Qian from Hangzhou Biogle Co., Ltd for rice transformation.

## References

1. Brinkman EK, Chen T, Amendola M, van Steensel B. 2014. Easy quantitative assessment of genome editing by sequence trace decomposition. Nucleic acids research 42(22): e168.

2. Goodwin S, McPherson JD, McCombie WR. 2016. Coming of age: ten years of next-generation sequencing technologies. Nature Reviews Genetics 17(6): 333–351.

3. Guell M, Yang L, Church GM. 2014. Genome editing assessment using CRISPR Genome Analyzer (CRISPR-GA). Bioinformatics 30(20): 2968–2970.

4. Li H, Durbin R. 2009. Fast and accurate short read alignment with Burrows-Wheeler transform. Bioinformatics 25(14): 1754–1760.

5. Lindsay H, Burger A, Biyong B, Felker A, Hess C, Zaugg J, Chiavacci E, Anders C, Jinek M, Mosimann C et al. 2016. CrispRVariants charts the mutation spectrum of genome engineering experiments. Nature biotechnology 34(7): 701–702.

6. Metzker ML. 2010. Sequencing technologies - the next generation. Nature Reviews Genetics 11 (1): 31.

7. Park J, Lim K, Kim JS, Bae S. 2016. Cas-analyzer: an online tool for assessing genome editing results using NGS data. Bioinformatics 33 (2): 286.

8. Pinello L, Canver MC, Hoban MD, Orkin SH, Kohn DB, Bauer DE, Yuan GC. 2016. Analyzing CRISPR genome-editing experiments with CRISPResso. Nature biotechnology 34(7): 695–697.

9. Rodriguez-Leal D, Lemmon ZH, Man J, Bartlett ME, Lippman ZB. 2017. Engineering Quantitative Trait Variation for Crop Improvement by Genome Editing. Cell 171(2):470–480.

10. Wang Y, Cheng X, Shan Q, Zhang Y, Liu J, Gao C, Qiu JL. 2014. Simultaneous editing of three homoeoalleles in hexaploid bread wheat confers heritable resistance to powdery mildew. Nature biotechnology 32(9): 947–951.

11. Xie X, Ma X, Zhu Q, Zeng D, Li G, Liu YG. 2017. CRISPR-GE: A Convenient Software Toolkit for CRISPR-Based Genome Editing. Molecular plant 10 (9): 1246–1249.

12. Xue LJ, Tsai CJ. 2015. AGEseq: Analysis of Genome Editing by Sequencing. Molecular plant 8(9): 1428–1430.

13. Zhang H, Zhang J, Wei P, Zhang B, Gou F, Feng Z, Mao Y, Yang L, Zhang H, Xu N et al. 2014. The CRISPR/Cas9 system produces specific and homozygous targeted gene editing in rice in one generation. Plant biotechnology journal 12(6): 797–807.

